# A bipartite interaction with the processivity clamp potentiates Pol IV-mediated TLS

**DOI:** 10.1101/2024.05.30.596738

**Authors:** Seungwoo Chang, Luisa Laureti, Elizabeth S. Thrall, Marguerite S Kay, Gaëlle Philippin, Slobodan Jergic, Vincent Pagès, Joseph J Loparo

## Abstract

Processivity clamps mediate polymerase switching for translesion synthesis (TLS). All three *E. coli* TLS polymerases interact with the β_2_ processivity clamp through a conserved clamp-binding motif (CBM), which is indispensable for TLS. Notably, Pol IV also makes a unique secondary contact with the clamp through non-CBM residues. However, the role of this “rim contact” in Pol IV-mediated TLS remains poorly understood. Here we show that the rim contact is critical for TLS past strong replication blocks. In *in vitro* reconstituted Pol IV-mediated TLS, ablating the rim contact compromises TLS past 3-methyl dA, a strong block, while barely affecting TLS past N^2^-furfuryl dG, a weak block. Similar observations are also made in *E. coli* cells bearing a single copy of these lesions in the genome. Within lesion-stalled replication forks, the rim interaction and ssDNA binding protein cooperatively poise Pol IV to better compete with Pol III for binding to a cleft through its CBM. We propose that this bipartite clamp interaction enables Pol IV to rapidly resolve lesion-stalled replication through TLS at the fork, which reduces damage induced mutagenesis.

## Introduction

Translesion synthesis (TLS) is a universal mechanism that cells use to resolve lesion-stalled DNA replication (1, 2). TLS is primarily mediated by TLS polymerases, which can efficiently synthesize past various template lesions that block replicative polymerases. TLS polymerases are largely error-prone, and their activities in cells are regulated at multiple steps ranging from transcription to protein-protein interactions (PPIs). Among these PPIs, interactions of TLS polymerases with processivity factors are indispensable for TLS in cells (3, 4).

The *E. coli* β_2_ clamp is an extensively characterized processivity factor that retains many of the structural and regulatory features of replicative processivity factors across species (5, 6). The *E. coli* β_2_ clamp is a homodimer, and each protomer has a binding cleft for conserved short peptides known as clamp binding motifs (CBMs). Clamp-interacting proteins (CLIPs) use these CBMs to interact with the β_2_ clamp (Fig 1A, top) (7–10). The DNA polymerases of *E. coli*, including all three TLS polymerases, Pol II, IV and V, have CBM(s). Mutating these CBMs abolishes DNA synthesis by these polymerases although these mutant polymerases retain wild-type polymerase activity, highlighting the importance of the CBM-clamp cleft interaction (4, 11).

**Fig 1.**
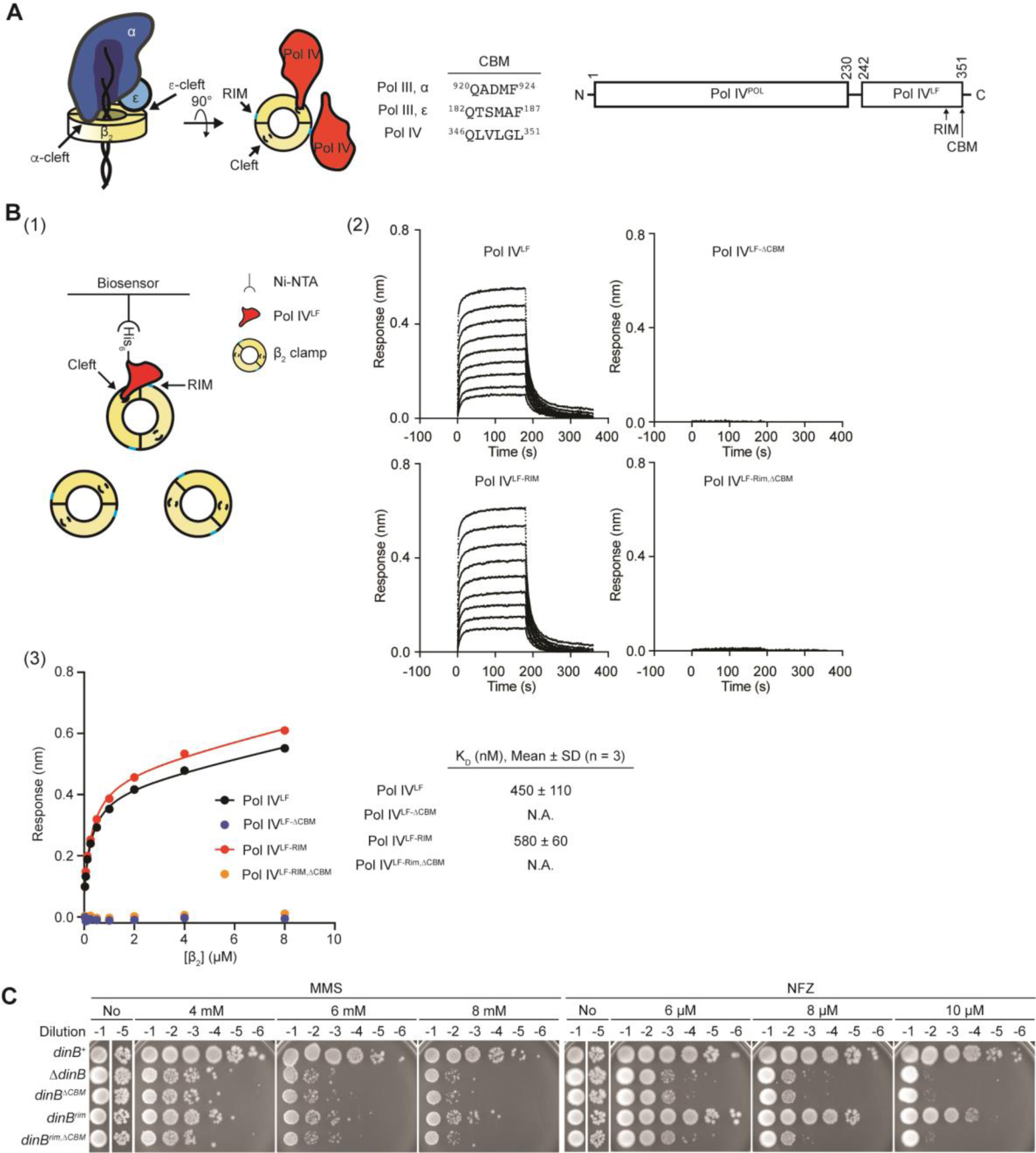
*dinB^rim^* is sensitized to damaging agents. A. (Left) Dual interaction of Pol III (αεθ) with the β_2_ clamp through CBMs of α and ε subunits and bipartite interaction of Pol IV^LF^ with the β_2_ clamp through the CBM and rim-interacting residues. (Middle) CBM sequences of Pol III and Pol IV. (Right) A domain structure of Pol IV. Pol IV^POL^, polymerase domain; Pol IV^LF^, little finger domain; RIM, rim-interacting residues (^303^VWP^305^); CBM, clamp binding motif (^346^QLVLGL^351^). B. Interaction of Pol IV^LF^ and β_2_ clamp. (1) Interaction between Pol IV^LF^ and β_2_ clamp was measured by biolayer interferometry. Indicated Pol IV^LF^ was immobilized to the NTA sensors through the N-terminal His_6_ tag and β_2_ clamp was titrated up to 8 μM. (2) Traces are responses in the presence of varying concentrations of β_2_ clamp. (3) Steady-state responses at given concentrations of β_2_ clamp were used to calculate equilibrium dissociation constants (K_D_) for the cleft. C. Mutating the rim-interacting residues of Pol IV sensitizes cells to MMS and NFZ. *dinB*, Pol IV; *dinB^rim^*, Pol IV with the rim-interacting residues mutated (^303^VWP^305^ to ^303^AGA^305^); *dinB^ΔCBM^*, Pol IV lacking the CBM; *dinB^rim, ΔCBM^*, Pol IV with both *dinB^rim^*and *dinB^ΔCBM^* mutations.

Switching between bacterial replicative (Pol III) and TLS polymerases for TLS is mediated by the β_2_ clamp. Underlying this polymerase switching is the dual interaction between Pol III and the β_2_ clamp (12). Pol III is a trimeric complex (αεθ) of the α polymerase subunit, the ε exonuclease subunit and the θ accessory subunit (13). Notably, the α and ε subunits have their own CBMs (CBMα and CBMε in Fig 1A, top) (11, 14), but CBMα interacts with the clamp with a much higher affinity than CBMε. During processive replication, the α and ε subunits simultaneously occupy both clefts of the β_2_ clamp (14), preventing the interaction of TLS polymerases with the β_2_ clamp (12, 14). However, upon lesion stalling, while Pol III remains associated with the β_2_ clamp through the α subunit, the cleft occupied by the ε subunit becomes more accessible to TLS polymerases, allowing a TLS polymerase to bind to the β_2_ clamp and mediate TLS past the lesion (12). This gatekeeping mechanism by the ε subunit is tuned to impose a limited kinetic window for TLS as weakening the clamp binding of the ε subunit increases TLS and vice versa (12). Persistent replication stalling due to failed TLS eventually leads to repriming of DNA replication. Therefore, the ε-cleft interaction plays a pivotal role in determining how stalled replication forks are resolved.

Despite the indispensable role of the β_2_ clamp in mediating TLS, the localization of Pol IV, and potentially other TLS polymerases, to lesion-stalled replication forks requires interaction with fork-associated ssDNA binding protein (SSB) (15, 16). While SSB is constitutively associated with replication forks, Pol IV only becomes enriched within lesion-stalled replication forks (17). Remodeling of fork-associated SSB upon lesion stalling drives this selective enrichment, enabling Pol IV to overcome the ε kinetic barrier (15, 18).

Interestingly, DNA polymerases also make non-cleft contacts with the β_2_ clamp that may have regulatory roles (8, 19). Such a non-cleft interaction was observed in a co-crystal structure of the β_2_ clamp and the little finger domain of Pol IV (Pol IV^LF^) (Fig 1A, right) (8). This additional contact located on the rim side of the β_2_ clamp, the “rim contact”, consists of a hydrophobic pocket formed by ^303^VWP^305^ of Pol IV^LF^ and L^98^ of the β_2_ clamp sticking into the pocket.

In this study, we demonstrate that the rim contact promotes Pol IV-mediated TLS by facilitating the interaction of Pol IV with the β_2_ clamp through the CBM. Binding of Pol IV to the β_2_ clamp through its CBM is competitively inhibited by the ε subunit of Pol III (12), and the rim contact enables Pol IV to better compete with the ε subunit of Pol III presumably by concentrating Pol IV in close proximity to the cleft. Dependence of Pol IV-mediated TLS on the rim contact becomes more critical for a strong replication block. Intriguingly, the rim contact is too weak for the interaction with the β_2_ clamp at cellular concentrations of Pol IV. We propose that Pol IV is enriched at stalled replication forks through interactions with SSB, which drives formation of the rim interaction, promoting TLS.

## Results

### Mutating the rim-interacting residues compromises Pol IV-mediated TLS in cells

In order to examine how the rim contact contributes to the interaction between Pol IV and the β_2_ clamp, we set up a biolayer interferometry (BLI)-based binding assay (20). In this assay, Pol IV^LF^ was immobilized onto Ni-NTA-coated BLI sensors through an N-terminal His_6_ epitope, and the interaction of Pol IV^LF^ with β_2_ clamp was detected by BLI (Fig 1B, left). Wild-type Pol IV^LF^ interacted with the β_2_ clamp with the equilibrium dissociation binding constant (K_D_) of ∼450 nM ((2) and (3) in Fig 1B), which is similar to the binding affinity measured by surface plasmon resonance with surface-immobilized β_2_ clamp(21). Mutating the rim-interacting residues (^303^VWP^305^ to AGA) within Pol IV (Pol IV^RIM^) only marginally weakened this interaction ((2) and (3) in Fig 1B), indicating that the rim contact is very weak. Furthermore, the rim interaction became measurable only in the presence of an intact CBM as deleting the CBM (Pol IV^ΔCBM^ and Pol IV^RIM,ΔCBM^) completely abolished the Pol IV-β_2_ clamp interaction ((2) and (3) in Fig 1B). These results suggest that while the rim interaction potentially promotes the CBM-β_2_ clamp interaction, the rim interaction by itself is too weak to be measured (K_D_ >> 20 μM).

Strikingly, however, the strain bearing the *dinB^rim^*mutation was highly sensitized to methyl methanesulfonate (MMS), a Pol IV cognate damaging agent, only slightly less than the *ΔdinB* strain (Fig 1C) (22, 23). In stark contrast, the *dinB^rim^* strain was only modestly sensitized to nitrofurazone (NFZ), another Pol IV cognate damaging agent (Fig 1C). The *dinB^rim^* strain retained a normal damage-induced SOS response including the induction of Pol IV (Fig S1A and B), ruling out the possibility that the increased sensitivity results from defective damage responses. In addition, the *dinB^ΔCBM^*, which is as sensitized as the *ΔdinB* mutant, was epistatic to *dinB^rim^* (Fig 1C), indicating that the increased sensitivity of the *dinB^rim^* is due to defective Pol IV-mediated TLS. These results suggest that the rim interaction promotes Pol IV-mediated TLS in cells and its contribution to TLS varies depending upon the lesion identity.

### The rim interaction is critical for TLS past 3meA

To understand the mechanistic basis of differential sensitivity of the *dinB^rim^* to NFZ and MMS, we reconstituted TLS *in vitro* within the *E. coli* replisome (12, 24). This reconstitution uses rolling circle DNA templates that bear on the leading strand template N^2^-furfuryl dG (N^2^-FFdG) or 3-deaza-methyl dA (3meA) (Fig 2A), which are structural analogs of major DNA lesions created in NFZ- and MMS-treated *E. coli* cells respectively (25, 26). In the absence of Pol IV, reconstituted replisomes only poorly replicated the N^2^-FFdG- and 3meA-containing templates (No Pol IV in Fig 2B), indicating that these are potent replication blocks for Pol III, the replicative polymerase (12, 15). Addition of Pol IV to the replication reactions promoted the synthesis of both leading and lagging strands with leading strand replication products accumulating at the resolution limit (RL) of the gel ((1) and (2) in Fig 2B) before Pol III-mediated replication is completely inhibited at higher concentrations of Pol IV (Fig S2B) (12, 15, 27, 28). Given that leading strand replication products accumulating at the RL result from more than 5 passages through blocking lesions without strand discontinuities such as nicks and gaps (Fig S2A), these results show that Pol IV can mediate TLS past these lesions within stalled replisomes (TLS at the fork) (12).

**Fig. 2.**
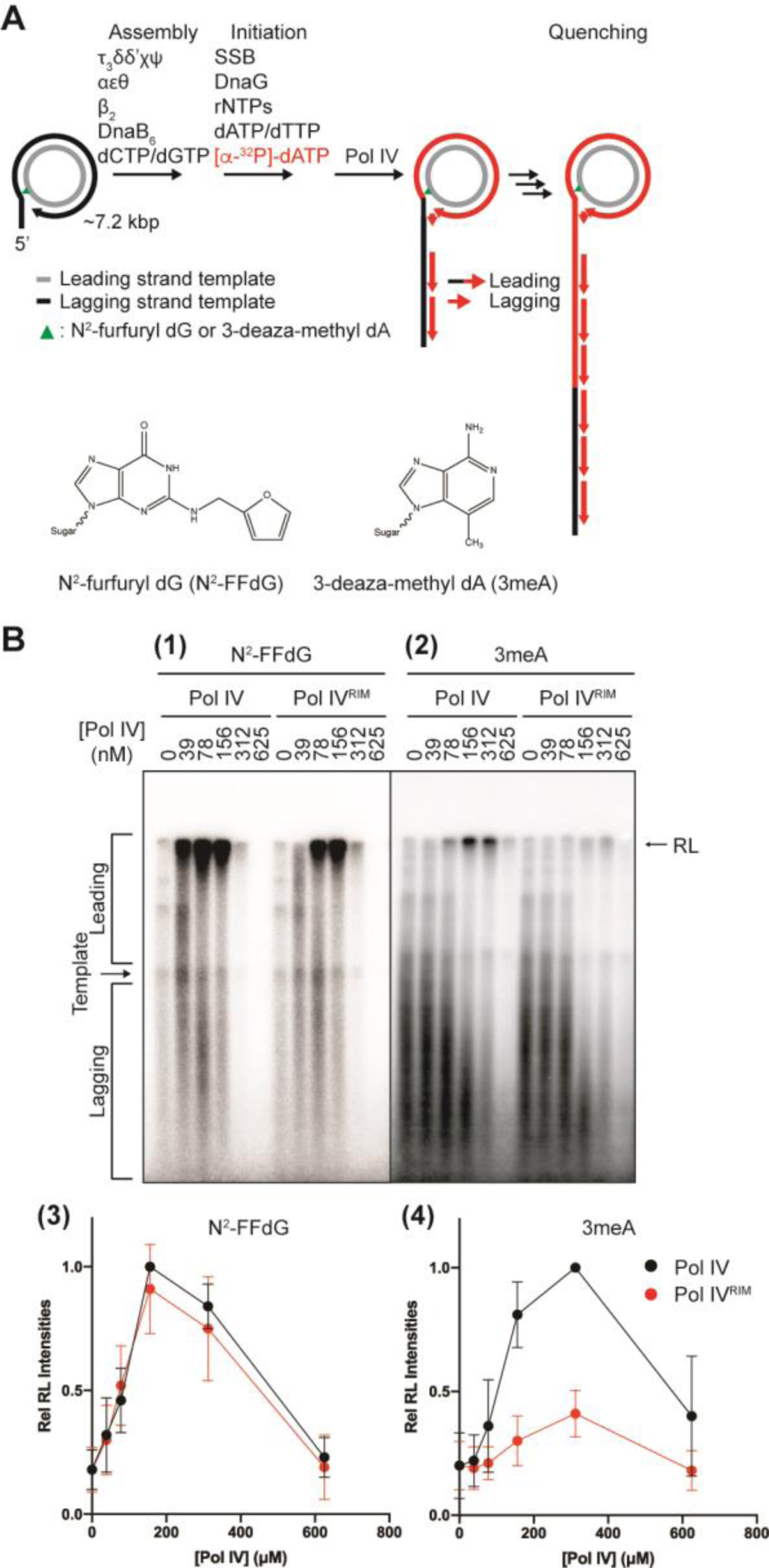
The rim interaction promotes polymerase switching within lesion-stalled replisome. A. A schematic of *in vitro* reconstituted Pol IV-mediated TLS. The inner circle (grey) is the leading strand template. Replication products are resolved in denaturing agarose gels and visualized by autoradiography as shown in Fig. 2B. B. Reconstitution of Pol IV-mediated TLS by Pol IV past N^2^-FFdG ((1) and (3)) and 3meA ((2) and (4)). Indicated amounts of Pol IV or Pol IV^RIM^ were added to the replication reactions on the lesion-containing rolling circle templates. RL, resolution limit of the gel, ∼45 kilonucleotides. Discrete bands between RL and template represent replication stalling and disappear in the presence of Pol IV. Rel. RL intensities, intensities of the replication products at RL relative to the highest intensity.

Our prior work with similar *in vitro* reconstitutions demonstrated that Pol IV is only briefly engaged in replication, primarily during the synthesis past a blocking lesion, while most of the replication is mediated by Pol III (12). When corrected for the replication-inhibitory effects of Pol IV (Fig S2B), maximum TLS past N^2^-FFdG and 3meA was achieved around 150 and 300 nM Pol IV respectively ((3) and (4) in Fig 2B). In addition, Pol IV-promoted replication of the N^2^-FFdG-containg template resulted in more leading strand replication products than that of the 3meA-containing template did (Fig 2B). Collectively, these results indicate that Pol IV-mediated TLS is less efficient within 3meA-stalled replication forks than it is within N^2^-FFdG-stalled replication forks.

Notably, Pol IV^RIM^ barely promoted the replication of the 3meA-containing template ((2) and (4) in Fig 2B) whereas Pol IV^RIM^ promoted replication of the N^2^-FFdG-containg template only slightly less efficiently as compared with Pol IV ((1) and (3) in Fig 2B). Together with Pol IV^ΔCBM^, which completely failing to promote TLS past N^2^-FFdG in our *in vitro* reconstitution (Fig S2C), these differential impacts of the rim interaction on *in vitro* reconstituted TLS correlate well with the sensitivity of the corresponding *dinB* strains to NFZ and MMS (Fig 1C). Therefore, we concluded that the rim interaction promotes TLS within lesion-stalled replisomes but plays a critical role only for strong replication blocks, such as 3meA.

### Cooperative interaction of Pol IV with SSB and β_2_ clamp

How the weak rim interaction robustly promotes the action of Pol IV within stalled replication forks remains unclear. Our prior work demonstrated that SSB highly enriches Pol IV at lesion-stalled replication forks, promoting formation of Pol IV-β_2_ clamp complexes (15). To examine whether the rim interaction contributes to the formation of Pol IV-β_2_ clamp complexes, we imaged individual Pol IV molecules localized at the fork in strains that express Pol IV-PAmCherry (Fig 3A, top) and SSB-mYPet as previously described (Fig 3A, bottom) (17). Briefly, we set the imaging conditions to selectively detect static Pol IV molecules, most of which represent Pol IV molecules forming complexes with SSB or the β_2_ clamp at lesion-stalled forks in MMS-treated cells (15). Shown in Fig 3B and C, radial distribution values (g(r)) represent the relative abundance of these complexes at a distance (radius) from the fork (radius = 0) relative to chance; random cellular localization is characterized by g(r) = 1 at all radius values, whereas g(r) > 1 indicates enrichment relative to chance. Deleting the CBM (Pol IV^ΔCBM^) caused a substantial reduction in the formation of Pol IV-β_2_ clamp complexes at the fork (g(r) = 5.7 ± 0.1, mean ± S.E.M here and thereafter) as compared with wild-type Pol IV (g(r) ∼ 8.0 ± 0.3). This reduction was slightly less than our previous measurement with Pol IV^RIM,ΔCBM^ (g(r) ∼ 5) (15), indicating that the CBM is the major contributor to the formation of Pol IV-β_2_ clamp complexes. Consistent with this idea, mutating only the rim interacting residues (Pol IV^RIM^) led to a small but discernable reduction (g(r) = 7.0 ± 0.1) (Fig 3B). However, when SSB-mediated fork enrichment of Pol IV was diminished by the *dinB^T120P^* mutation(15), the *dinB^rim^* mutation (Pol IV^T120P,RIM^) did not reduce the Pol IV-β_2_ clamp interaction (g(r) = 3.3 ± 0.1) compared with our previous measurement with Pol IV^T120P^ (g(r) ∼3) (15); combining the T120P mutation with deletion of the CBM (Pol IV^T120P, ΔCBM^) completely ablated complex formation (g(r) = 0.47 ± 0.01) (Fig 3C). Collectively, these results suggest that the rim interaction is utilized within lesion-stalled replication forks but only when Pol IV is highly enriched by SSB.

**Fig. 3.**
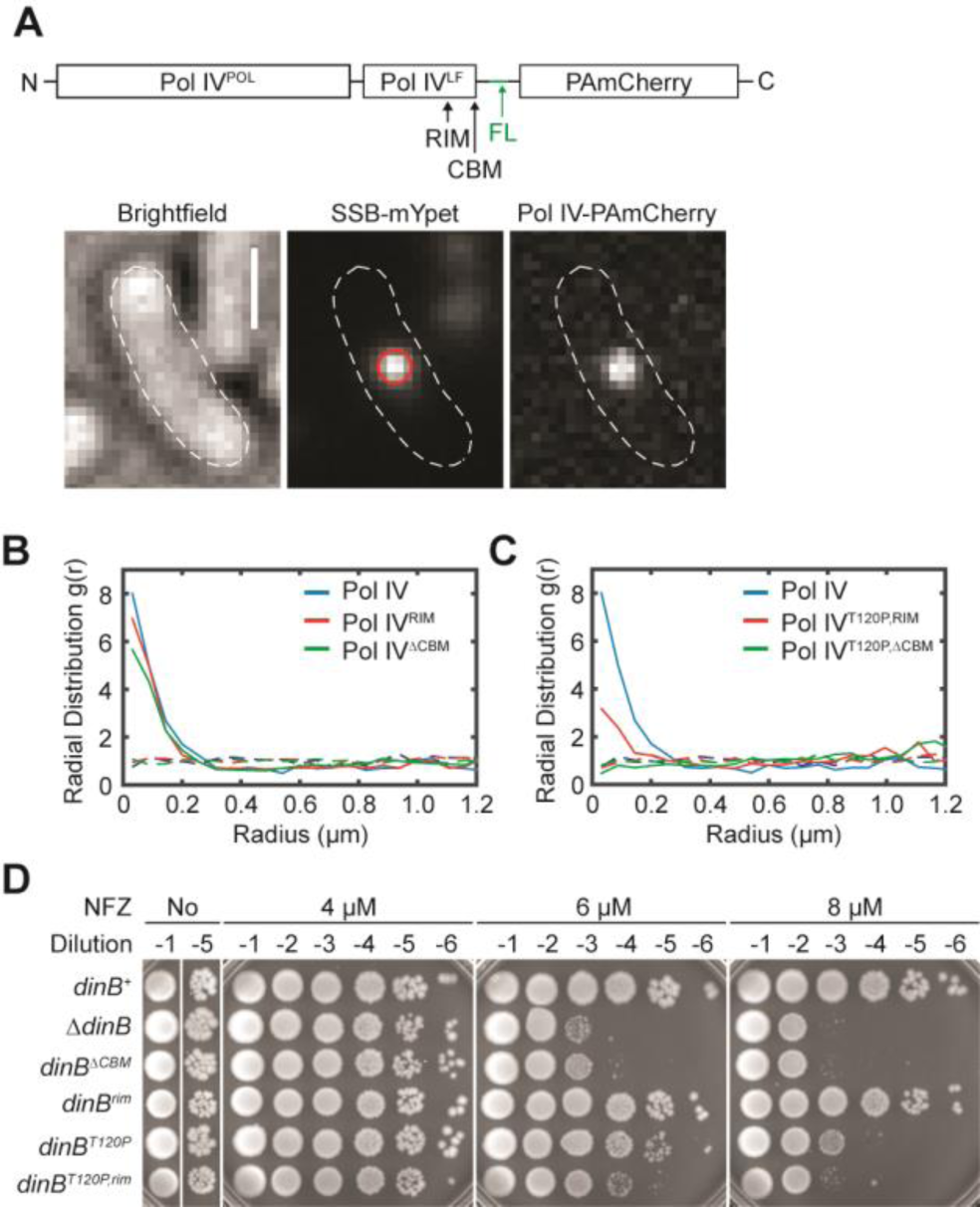
Cooperative interaction of Pol IV with SSB and β_2_ clamp at the fork. A. A schematic of Pol IV-PAmCherry fusion (Top) and representative micrographs of an imaging strain (Bottom). (Top) Pol IV-PAmCherry. Pol IV^POL^, polymerase domain; Pol IV^LF^, little finger domain; RIM, rim-interacting residues, (^303^VWP^305^); CBM, clamp binding motif (^346^QLVLGL^351^); FL, flexible linker (G_4_S)_4_. (Bottom) Transmitted white light micrograph of single *E. coli* cell with cell outline and 1 μM scalebar (Left), fluorescence micrograph of single SSB-mYPet focus with overlaid centroid (Middle), and fluorescence micrograph of single Pol IV-PAmCherry molecule (Right). B. Radial distribution function g(r) analysis of Pol IV in cells treated with 100 mM MMS for 20 min for (B) Pol IV^WT^ (*N* = 1,436), Pol IV^RIM^ (*N* = 3,905), and Pol IV^ΔCBM^ (*N* = 3,753) and (C) Pol IV^WT^ (*N* = 1,436), Pol IV^T120P,RIM^ (*N* = 1,881), and Pol IV^T120P, ΔCBM^ (*N* = 2,958). N, number of static Pol IV molecules included in the analysis. D. Sensitivity to NFZ of the indicated strains.

In contrast to its high enrichment in MMS-treated cells, Pol IV is only slightly enriched at the fork in NFZ-treated cells (17), and the *dinB^T120P^* strain is only modestly sensitized to NFZ (Fig 3D) (15), suggesting that relatively low enrichment is sufficient for Pol IV to mediate TLS past weak replication blocks such as structural analogs of N^2^-FFdG. Notably, even though ablating the interaction of Pol IV with either SSB (*dinB^T120P^*) or the rim (*dinB^rim^*) only modestly sensitized cells to NFZ, ablating both the SSB and the rim interactions (*dinB^T120P,rim^*) severely sensitized cells to NFZ (Fig 3D). These results suggest that the rim promotes TLS past weak replication block as well, but its impact is much smaller. We propose that although the rim contact by itself is intrinsically very weak (Fig 1B), SSB-mediated enrichment of Pol IV within stalled replication forks drives the rim interaction, further concentrating Pol IV near the β_2_ clamp.

### The rim interaction facilitates the CBM-cleft interaction

Interaction of Pol IV with β_2_ clamp through its CBM is indispensable for TLS (Fig 1C and S2C) (15). However, this interaction is competitively inhibited by the ε subunit of Pol III, which acts as a gatekeeper at the replication fork (12). To see whether the rim interaction helps Pol IV overcome this gatekeeping mechanism, we examined the effect of strengthening the interaction between the ε subunit and the β_2_ clamp on Pol IV-mediated TLS in cells. Consistent with our prior work (12), strengthening the interaction with the *ε_L_* mutation in *dnaQ* (14) (Fig 4A, see also Fig 1A), which enhances the gatekeeping mechanism (12), modestly sensitized the cells to NFZ (Fig 4B). This sensitization reflects suppressed Pol IV-mediated TLS because *dinB^ΔCBM^* was epistatic to *dnaQ(ε_L_)* (Fig 4B). Notably, *dinB^rim^* with the *dnaQ(ε_L_)* background (*dnaQ(ε_L_) dinB^rim^*) was highly sensitized to NFZ even though *dinB^rim^* with *dnaQ^+^* background (*dnaQ^+^ dinB^rim^*) was only modestly sensitized (Fig 4B). These results suggest that the rim interaction enables Pol IV to better compete with the ε subunit.

**Fig. 4.**
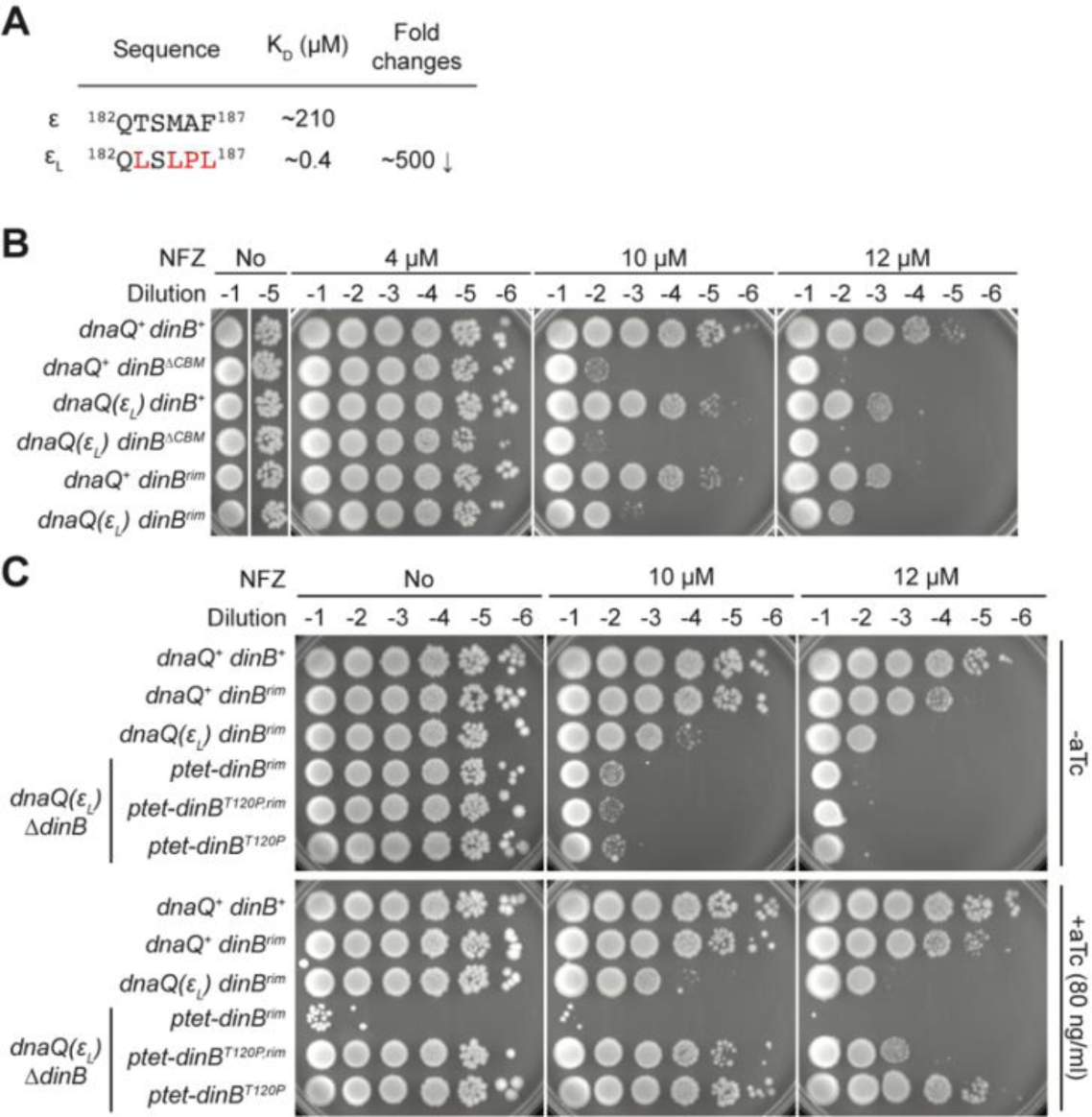
The rim interaction potentiates the CBM-cleft interaction. A. A CBM mutation in the ε subunit that strengthens the ε-clamp interaction. B. Sensitivity to NFZ of the indicated strains. C. Elevating the expression of Pol IV^RIM^ in *dnaQ(ε_L_) ΔdinB* normalizes tolerance to NFZ. Indicated Pol IV variants were ectopically expressed from a Tet-inducible expression cassette engineered in the *E. coli* genome.

Next, we tested whether loss of the rim interaction could be compensated for by elevating the expression of Pol IV^RIM^ in cells above its native levels. However, like Pol IV, elevated levels of Pol IV^RIM^ inhibited Pol III-mediated processive replication (Fig S2B), resulting in lethality (No NFZ in Fig 4C and S4A) (28, 29). Therefore, we leveraged the *dinB^T120P^* background, which substantially attenuates such lethality (No NFZ in Fig 4C and S4A)(15, 22). Elevating the expression level of Pol IV^T120P,RIM^ in *dnaQ(ε_L_) ΔdinB* made the cells more tolerant to NFZ (compare -aTc and +aTc in Fig 4C), even more than *dnaQ(ε_L_) dinB^rim^* (compare *dnaQ(ε_L_) dinB^rim^* and *dnaQ(ε_L_) ΔdinB ptet-dinB^T120P,rim^* in Fig 4C). These results indicate that the role of the rim interaction is concentrating Pol IV near the β_2_ clamp. In addition, elevating the expression level of Pol IV^T120P^ in *dnaQ(ε_L_) ΔdinB* made the strain even more tolerant to NFZ compared with that of Pol IV^T120P,RIM^ (Fig 4C). These results support the model that Pol IV is sequentially concentrated near the β_2_ clamp by its interactions with SSB and the rim.

### The impact of the rim interaction on the pathway choice of lesion-stalled replication

To directly examine whether the rim interaction promotes TLS in cells, we employed a genetic assay by which resolution of lesion-stalled replication through TLS or damage avoidance (DA) is quantified (30). Briefly, a single replication blocking lesion is integrated into the leading strand template of the *E. coli* genome (Fig 5A, left), and the resolution pathway choice in individual cells is determined by the color of the resulting colonies on X-gal plates (Fig 5A, right). In the *dinB^+^* strain, 3meA-stalled replication was primarily resolved by TLS (∼80%) while only a small fraction (∼20%) was through DA, which reflects repriming and subsequent homology-directed gap repair (Fig 5B). Consistent with Pol IV being the major TLS polymerase for 3meA, deletion of Pol IV (*ΔdinB*) resulted in a substantial increase in DA (∼95%) (Fig 5C). In the *dinB^rim^* mutant, resolution through TLS was reduced by ∼20% with a concomitant increase in resolution through DA, indicating that the rim interaction indeed promotes Pol IV-mediated TLS in cells. This reduction is smaller than the impact of the *dinB^rim^* mutation on *in vitro* reconstituted TLS past 3meA (Fig 2B). However, it should be noted that our rolling circle-based assay amplifies reductions in TLS because the reduction accumulates over multiple passages through the blocking lesion. As our cell-based assay is single passage, both the in cell and the *in vitro* observations are consistent. In the *dinB^+^* strain, N^2^-FFdG-stalled replication was also primarily resolved through TLS (∼60%) (Fig 5C) (15). In stark contrast to TLS past 3meA, however, TLS past N^2^-FFdG was not reduced in the *dinB^rim^* mutant (Fig 5B), which is consistent with our *in vitro* results (Fig 2B). Additionally, in the *ΔdinB* background, in which TLS can be inefficiently mediated by Pol III (15), TLS-mediated resolution of 3meA-stalled replication was only ∼7 % whereas that of N^2^-FFdG-stalled replication was ∼40%, indicating that 3meA is a much stronger replication block for Pol III than N^2^-FFdG (Fig 5C). Collectively, these results demonstrate that the rim interaction indeed promotes TLS-mediated resolution of lesion-stalled replication in cells and plays a more pronounced role for a stronger replication block.

**Fig. 5.**
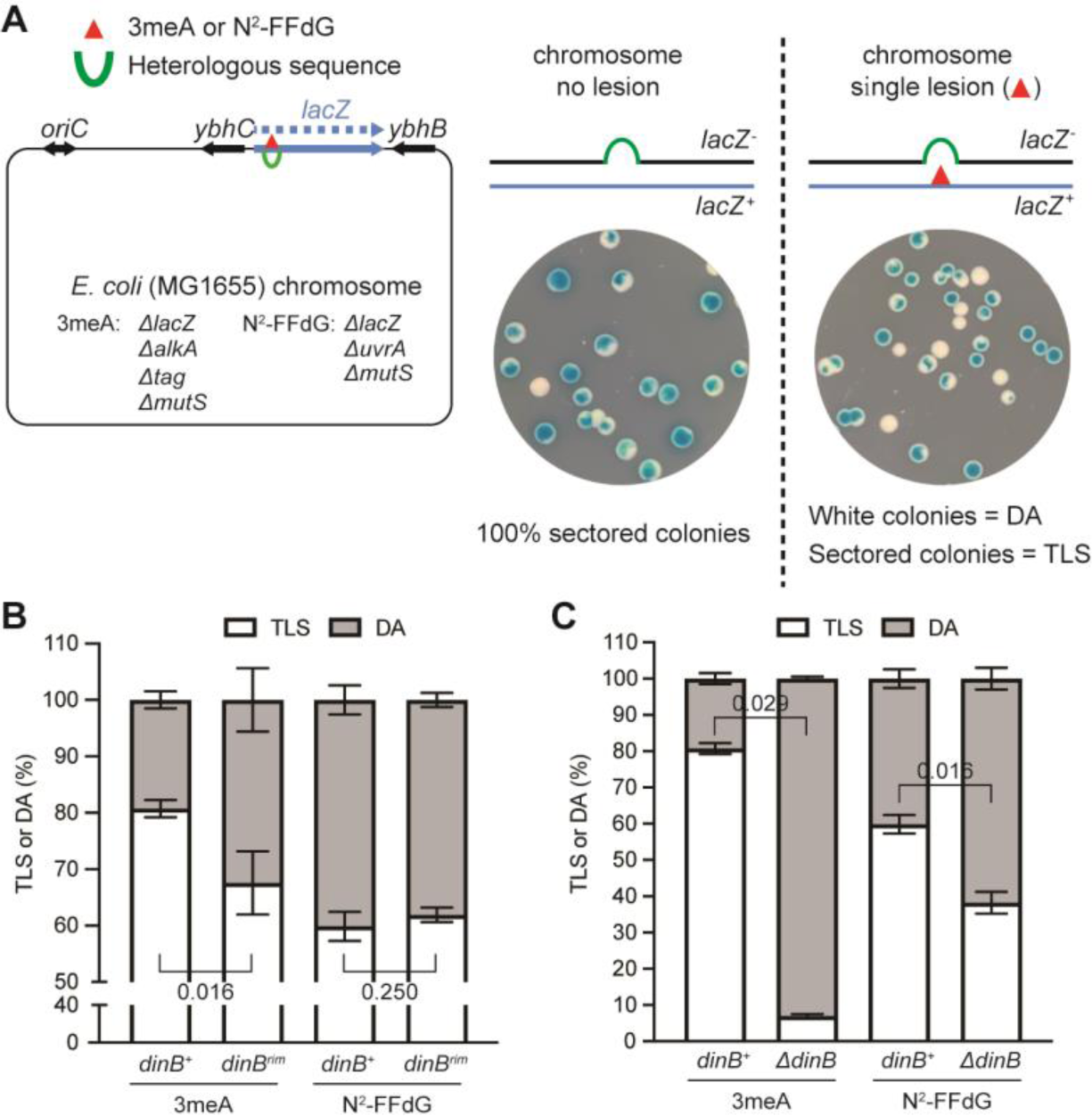
Pathway choice of lesion-stalled replication in cells. A. Quantification of pathway choice between TLS and DA in cells. (Left) The assay strain is based on *E. coli* MG1655 and the genome was engineered as shown in the diagram. To prevent the genome-incorporated lesions from being removed by repair pathways, base excision (*alkA* and *tag*) or nucleotide excision (*uvrA*) repair was inactivated together with the inactivation of the mismatch repair (*mutS*). Right, On X-gal plates, resolution of a lesion-stalled replication through DA in a cell results in the formation of a white colony whereas resolution through TLS results in the formation of a blue-sectored colony. Note that functional LacZ is expressed only when stalled replication is resolved by TLS. In this study, blocking lesions were introduced into the leading strand template in the *E. coli* genome. All the results presented here are based on at least 3 measurements with independently prepared materials. B. Pathway choice of lesion-stalled replication in *dinB^+^*and *dinB^rim^*. p values for the indicated comparison pairs were calculated with Mann-Whitney and Wilcoxon matched pairs test. C. Contribution of Pol IV to TLS-mediated resolution of lesion-stalled replication.

## Discussion

### Hierarchical protein-protein interactions for utilization of TLS polymerases

The rim interaction was initially observed in a co-crystal structure of the little finger domain of Pol IV (Pol IV^LF^) and the β_2_ clamp (8). In the structure, the C-terminal end of the Pol IV^LF^, which contains the CBM of Pol IV, assumes a stretched conformation in which Pol IV^LF^ makes simultaneous contacts with both a cleft and the adjacent rim. However, the interaction through the rim contact is very weak (Fig 1B), and it is possible that the rim interaction is stabilized by the CBM-β_2_ interaction under crystallization conditions. How does such a weak interaction have physiological relevance? Pol IV is highly enriched at lesion-stalled replication forks through the interaction with replisome-associated SSB (15, 17, 18). Therefore, the concentration of Pol IV near the β_2_ clamp could be high enough to drive the weak rim interaction. This transient but frequent interaction at the rim potentially poises Pol IV to more efficiently bind to the adjacent cleft, which is competitively inhibited by Pol III and possibly other clamp interacting proteins (12, 15). This competitive advantage imparted by the rim interaction becomes more important for lesions that are difficult to bypass.

As inferred from the co-crystal structure, a strong rim interaction would keep the polymerase domain away from the Primer/Template junction (8), preventing Pol IV-mediated synthesis. Indeed, *dinB^rim^*mutations tested by others (^303^VWP^305^ to AGA or SGA) slightly increased β_2_ clamp-supported primer extension by Pol IV (21, 22, 29). Moreover, a strong rim interaction would allow Pol IV to occupy the cleft more potently, inhibiting Pol III-mediated processive replication (12, 27, 31). Therefore, in conjunction with the selective enrichment of Pol IV at stalled forks by SSB (12, 15), a weak rim interaction promotes Pol IV-mediated TLS without perturbing replication.

### The role for Pol IV-mediated TLS in damage-induced mutagenesis

MMS-induced mutagenesis serves as a good model for how different TLS polymerases are utilized during the resolution of lesion-stalled replication. 3-methyladenine, a major methyl adduct in MMS-treated cells, is a cognate lesion of Pol IV, yet MMS-induced mutagenesis requires *umuDC*, the gene encoding Pol V (32). This Pol V-dependent mutagenesis is highly increased upon deletion of *dinB* (23). As with other leading strand lesions, methyl adduct-stalled replisomes can reprime DNA replication downstream of the blocking lesion(15), which leaves a ssDNA gap (33, 34). These gaps can be filled in by post-replicative synthesis involving Pol V, a highly mutagenic TLS polymerase (35). Our prior work demonstrated that Pol IV can mediate TLS within lesion-stalled replisomes (TLS at the fork), preventing replisomes from repriming (12). In MMS-treated *ΔdinB*, it is likely that lesion-stalled replisomes more frequently reprime, resulting in elevated mutagenesis (22, 23). In *dinB^rim^*, the MMS-induced SOS response (Fig S1C), which reflects the generation of ssDNA gaps, and MMS-induced mutagenesis are both elevated, but less so compared to *ΔdinB* (22). These observations are consistent with the rim interaction promoting Pol IV-mediated TLS at the fork in cells which suppresses damage-induced mutagenesis.

In light of damage-induced mutagenesis, what is the benefit of having this ancillary protein-protein interaction? Our results suggest that the rim interaction allows for more efficient TLS at the fork without high induction of Pol IV, which can inhibit DNA replication and repriming, thus preventing rapid resolution of lesion-stalled replication.

### Possible roles for ancillary clamp interactions of DNA polymerases

All DNA polymerases of *E. coli*, except for Pol I, contain conserved CBMs (7), which are indispensable for their functions in cells (4, 36). However, it is possible that like Pol IV, these polymerases also make secondary contacts with the β_2_ clamp that modulate the CBM-β_2_ clamp interaction. For example, Pol III, the replicative DNA polymerase, makes additional contacts with the β_2_ clamp and ablating these interactions impair the action of Pol III (37–39). Given the impact of the secondary clamp interaction of Pol IV on damage-induced mutagenesis, it would be interesting to discover such auxiliary clamp interactions for other TLS polymerases.

## Acknowledgments

We thank K. Arnett (Center for Macromolecular Interactions, Harvard Medical School) for assistance with the BLI experiments, N. Dixon (University of Wollongong) for feedback on our manuscript and D. Li (University of Rhode Island) for providing the N*^2^-*FFdG-containing oligomer. This work was supported by National Institutes of Health grants R01 GM114065 (to J.J.L.) and F32 GM113516 (to E.S.T.), The William F. Milton Fund (to S.C.), Fondation pour la Recherche Médicale Equipe FRM-EQU201903007797 (to V.P.).

---

**Fig. S1.**
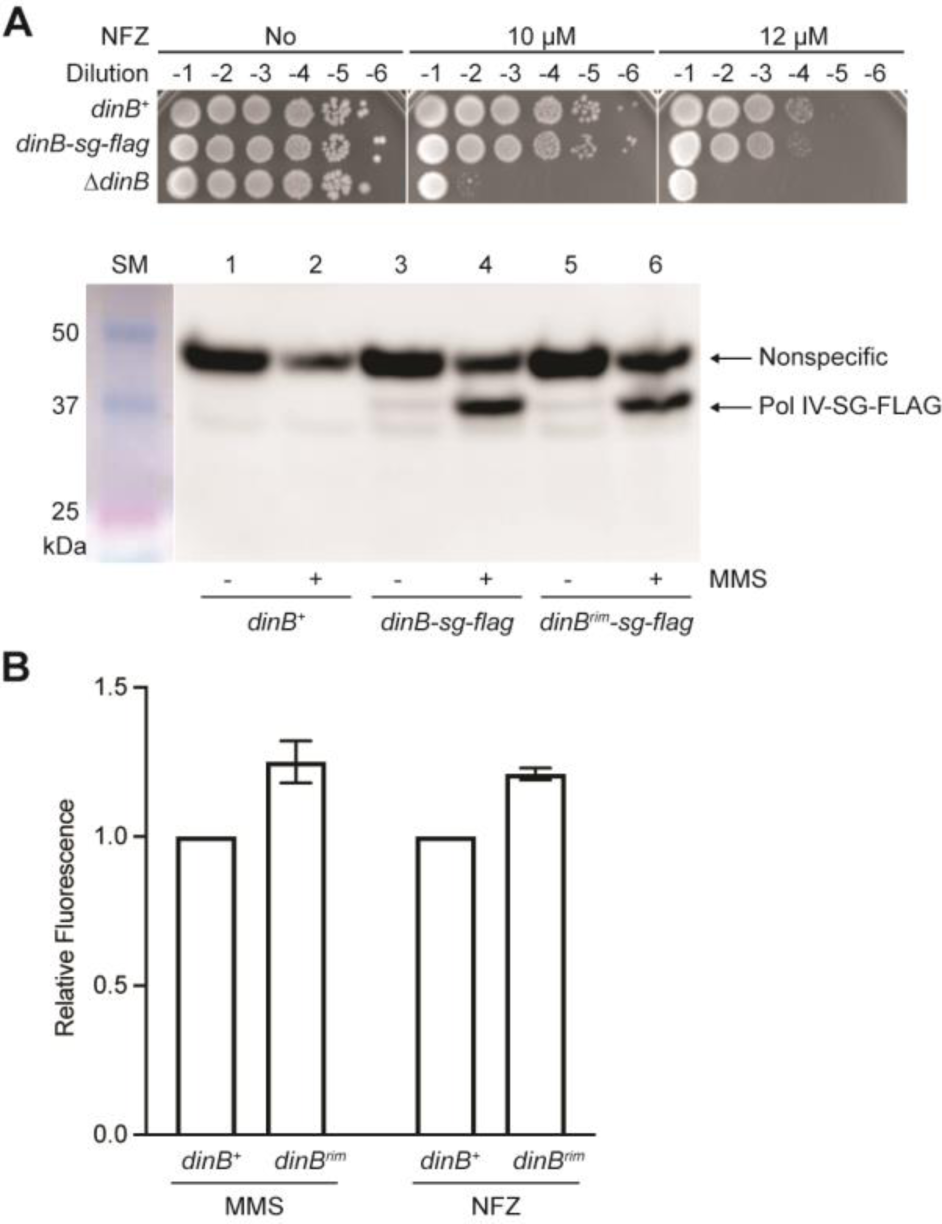
A. Expression of Pol IV and Pol IV^RIM^ in cells. To probe Pol IV and Pol IV^RIM^ in cells by Western blotting, a sequence encoding a FLAG epitope was added to the 3ʹ end of the *dinB* gene in the *E. coli* chromosome (*dinB-flag*). (Top) *dinB-flag* retained wild-type tolerance to NFZ. (Bottom) Indicated strains were treated with mock or MMS (7.5 mM) for 1 hour and cell lysates were resolved by SDS-PAGE and probed with an anti-FLAG antibody. SM, size markers. B. Damage-induced SOS responses in *dinB^+^* and *dinB^rim^*. The GFP fluorescence-based SOS reporter strain bearing *dinB^+^* or *dinB^rim^* was treated with 7.5 mM MMS or 60 μM NFZ and the expression of GFP in individual cells was measured by flow cytometry. Shown are means and SDs of three independent measurements.

**Fig. S2.**
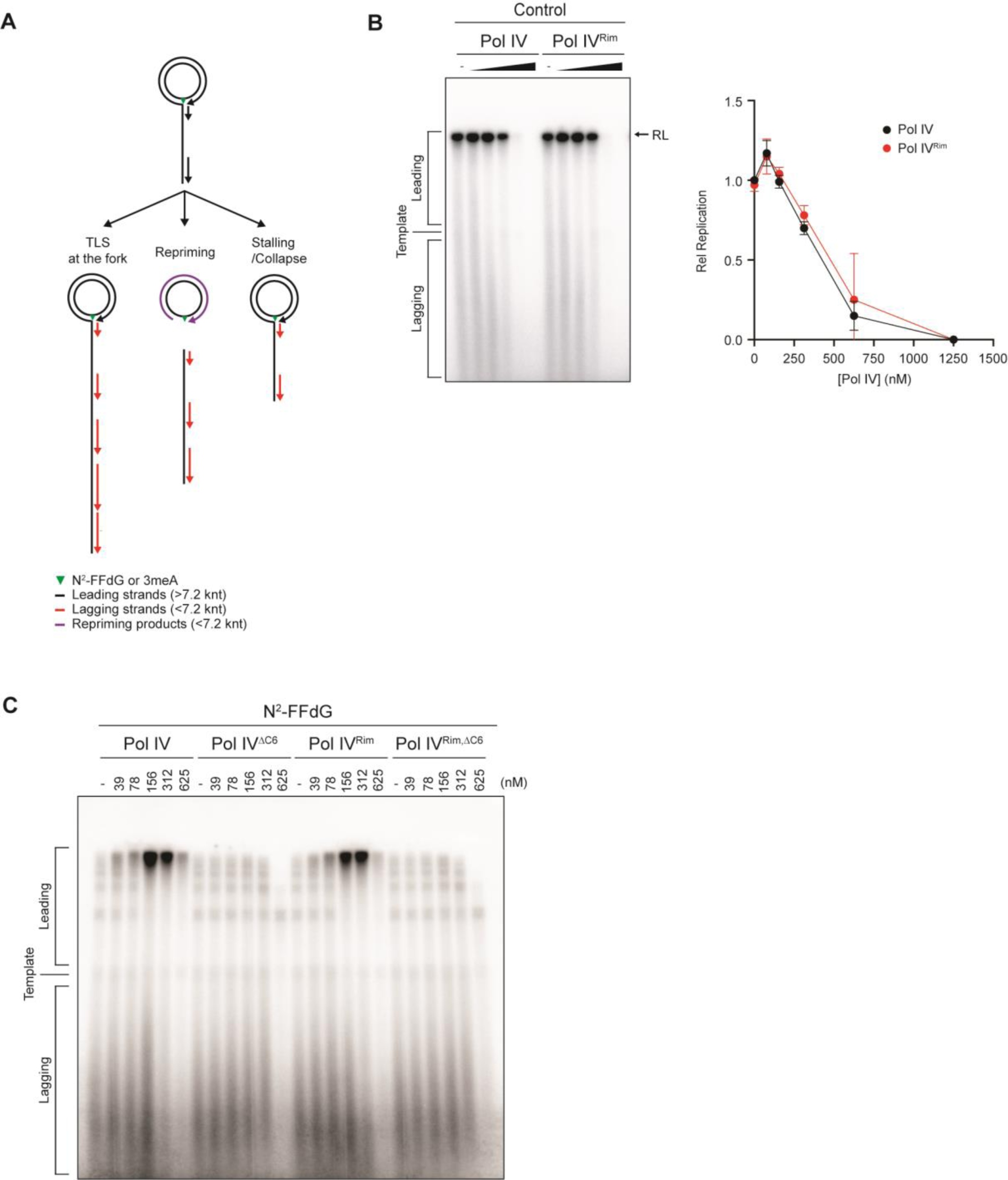
A. Fates of replisomes and associated replication products. TLS at the fork results in the leading strand replication products longer than the template. Repriming results in the leading strand replication products shorter than the template. Stalling/Collapse results in the leading strand replication products between RL and template. B. Inhibition of Pol III-mediated replication by Pol IV is barely reduced by the *dinB^rim^* mutation. Indicated amounts of either Pol IV or Pol IV^RIM^ were added to the replication reactions on a lesion-free rolling circle template. C. Interaction of Pol IV with β_2_ clamp through CBM is indispensable for TLS. Indicated amounts of Pol IV, Pol IV^ΔCBM^, Pol IV^RIM^ or Pol IV ^RIM,ΔCBM^ were added to the replication reactions on the N^2^-FFdG-containing rolling circle template.

**Fig. S3.**
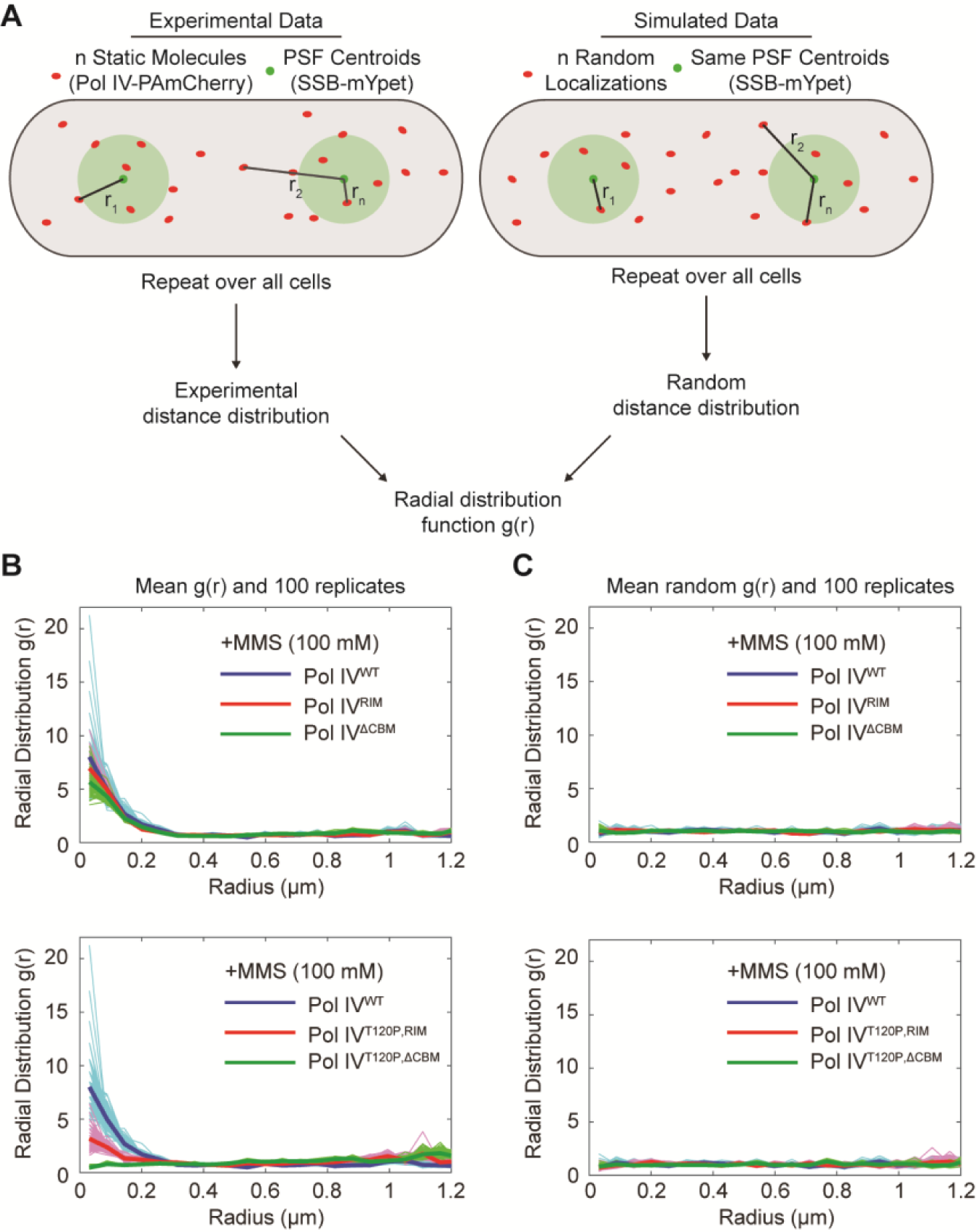
A. Cartoon of radial distribution function analysis. Distances between static Pol IV-PAmCherry molecules and the nearest SSB-mYPet focus are calculated for (Left) experimental Pol IV localizations and (Right) randomly simulated Pol IV localizations. Distances are aggregated for all cells and the radial distribution function *g(r)* is calculated by normalizing the experimental distribution by the random distribution. This procedure is repeated using 100 independent random distance distributions and the resulting 100 *g(r)* curves are averaged to give the mean *g(r)* curve. A separate random distance distribution is generated and normalized following the same approach to give a mean random *g*(*r*) curve. B. The 100 individual Pol IV-SSB radial distribution function curves (thin lines) and the mean *g*(*r*) curve (thick lines) for Pol IV^WT^ and mutants in MMS-treated cells. (Top) Pol IV^WT^, Pol IV^RIM^, and Pol IV^ΔCBM^. (Bottom) Pol IV^WT^, Pol IV^T120P,RIM^, and Pol IV ^T120P,ΔCBM^. C. As in B, but for the random *g(r)* curves.

**Fig. S4.**
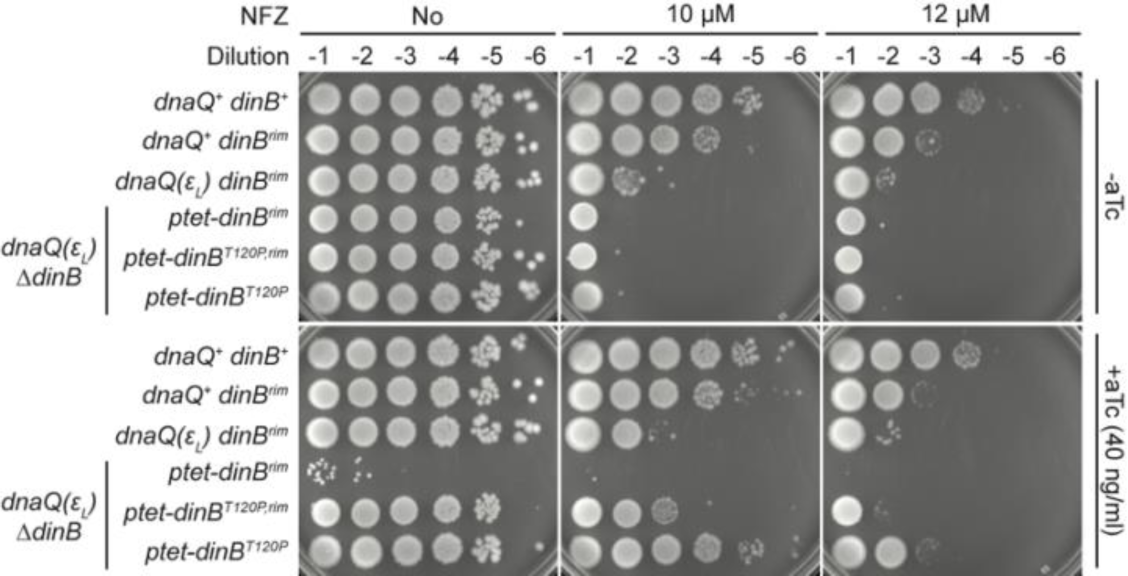
Elevating the expression of Pol IV^RIM^ in *dnaQ(ε_L_) ΔdinB* normalizes tolerance to NFZ. Indicated Pol IV variants were ectopically expressed from a Tet-inducible expression cassette engineered in the *E. coli* genome.

**Fig. S5.**
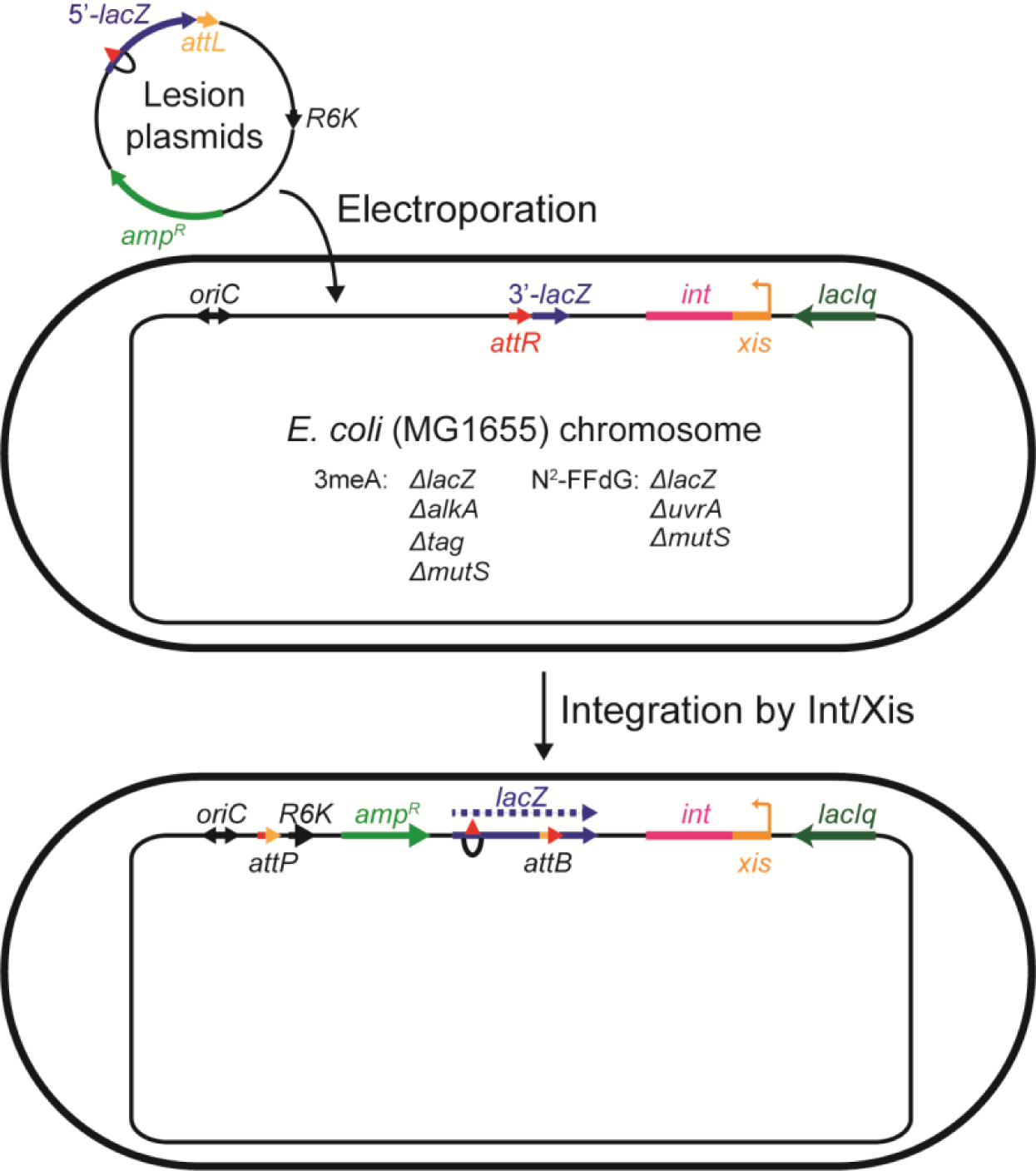
Integration of a replication blocking lesion into the *E. coli* genome. The assay strains with the indicated modifications are transformed with lesion-containing non-replicative plasmids. These plasmids are incorporated into the *E. coli* genome through the recombination between *attL* (plasmid) and *attR* (genome) by Integrase (Int) and Excisionase (Xis). The resulting integrants become resistant to ampicillin and gain a single copy of the coding sequence for LacZ in either the leading or lagging strand.

**Table 1.**
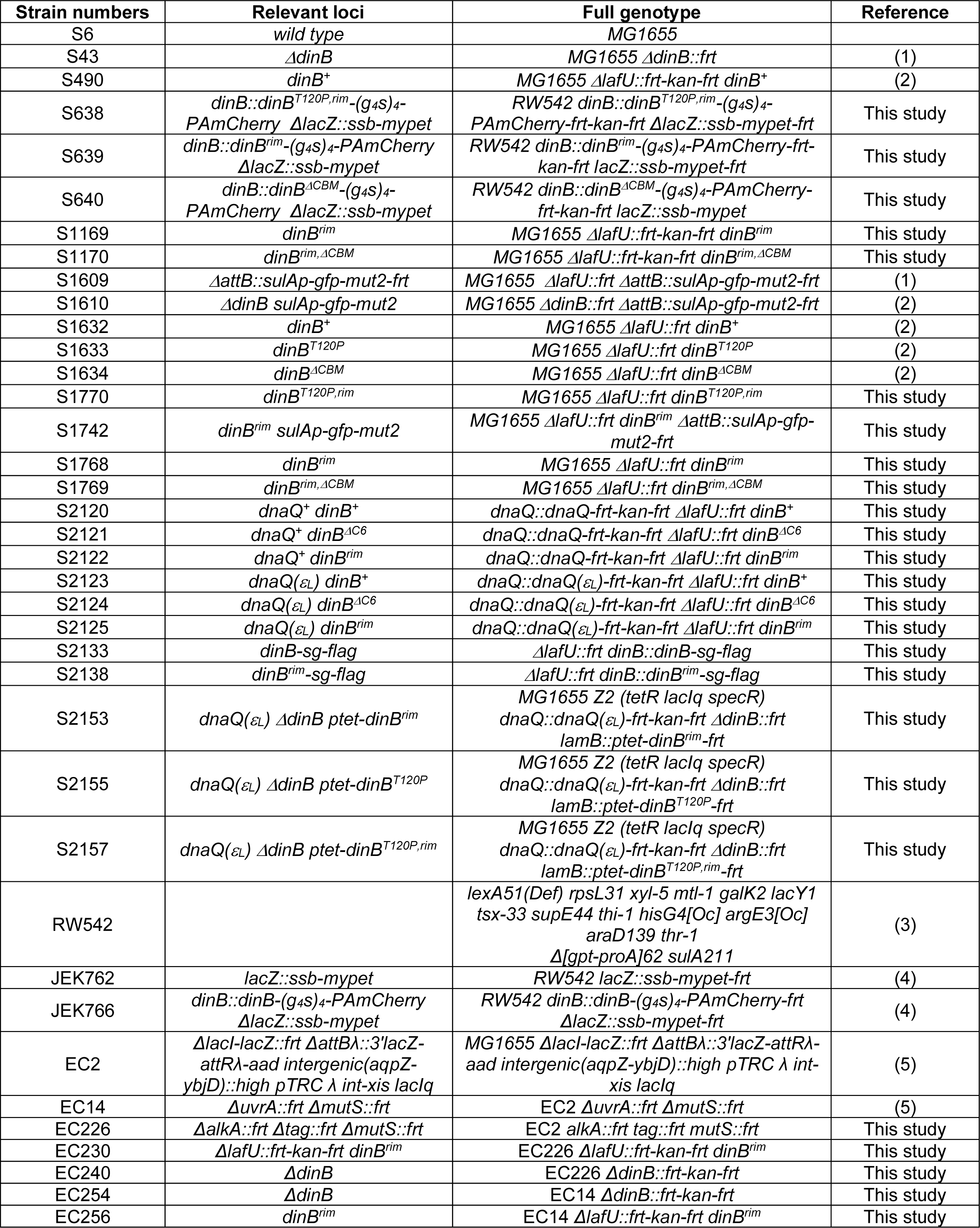
Strains used in this study.

## Materials and Methods

### Chemicals

Anhydrotetracycline (aTc, TaKaRa, 631310), ATPγS (Abcam, AB138911), dNTPs (New England Biolabs, N0446S), N,N-Dimethyl formamide anhydrous (Sigma, 227056), Formaldehyde (Sigma, F8775), Methyl methanesulfonate (MMS, Sigma, 129925), and Nitrofurazone (NFZ, Fluka, PHR1196) and rNTPs (New England Biolabs, N0450S).

### Construction of strains

The *E. coli* strains used in this study are all based on MG1655 and were created as described previously (1, 2). Briefly, recombineering strains bearing pSIM6 (6) were transformed with PCR-amplified dsDNA fragments containing an antibiotic selection marker and a desired modification(s). In some cases, modified genomic loci were transferred into recipient strains by P1 transduction. From antibiotic-selected transformants, genomic regions containing the desired modifications were PCR amplified and sequenced. To flip out selection markers, cells were transformed with pCP20, which expresses a flippase, and the removal of a selection marker was confirmed by PCR. Construction of the parental strains for the pathway choice assay were described previously (7, 8).

### Proteins

*E. coli* replication proteins were purified as described previously: Pol III clamp loader complex (τ_3_δδ’χψ) (9), Pol III core (αεθ) (10), DnaB helicase (11), DnaG primase (12), β_2_ sliding clamp (13) and SSB (14). Full-length wild-type Pol IV and its variants were expressed in BL21(DE3) *dinB::frt-kan-frt* and purified as described previously (15). PK-His6-tagged Pol IV^LF^ (Pol IV^LF^: a.a. 243 - 351 of Pol IV with MGLRRASVHHHHHHSSGHIEGRHM appended to the N-terminus) and its variants were expressed in Rosetta2 and purified using Ni-NTA resins. Eluates from Ni-NTA resins were further purified by size exclusion chromatography with a Superose 6 increase 10/300 GL column (Cytiva). Purified PK-His6-Pol IV^LF^ and its variants were stored in a storage buffer (50 mM Hepes-NaOH, pH8/100 mM NaCl/5% glycerol/10 mM 2-mercaptoethanol).

### Sensitivity to DNA damaging agents

Overnight cultures grown in LB at 30 °C with aeration were 10-fold serially diluted (from OD_600_ = 0.1) in LB and spotted onto LB agar plates containing NFZ or MMS, and anhydrous tetracycline (aTc) when overexpression of proteins of interest was needed.

### Western blotting

The expression levels of Pol IV-FLAG and Pol IV^RIM^-FLAG were measured by Western blot as described previously (2). Overnight cultures grown in LB at 37 °C with aeration were diluted into fresh LB to OD_600_ ≈ 0.1 and incubated at 37 °C with aeration until OD_600_ reached about 0.5. Then, MMS (final 7.5 mM) or a mock was added to the culture and cultures were further incubated at 37 °C with aeration for 1 hour before cells were harvested. Cell lysates were prepared by lysing cells in B-PER^TM^ (Thermo Scientific^TM^) supplemented with Benzonase nuclease (Novagen, #70746) and EDTA-free protease inhibitor cocktail (Roche, #04693159001) and, based on OD_600_ at the time of harvest, an equal number of cells were loaded and run on an SDS-PAGE gel (Bio-Rad #4561086: 4-15 Mini-PROTEAN TGX) along with Precision Plus Protein Dual Color Standards (BIO-RAD, 1610374). Resolved proteins on an SDS-PAGE gel were transferred to a polyvinylidene difluoride membrane (PerkinElmer, #NEF1002001PK: PolyScreen PVDF Hybridization Transfer Membrane). After blocked in PBST containing 5% skim milk, the membrane was probed with a rabbit anti-FLAG polyclonal antibody raised against the antigen Ac-C(dPEG4)DYKDDDDK-OH (a gift of Johannes Walter, Harvard Medical School). A goat anti-rabbit IgG-HRP antibody (Jackson ImmunoResearch, #111-035-003) was used as a secondary antibody. The membrane and chemiluminescent signals were visualized using an Amersham Imager 600 with HyGLO chemiluminescent HRP antibody detection reagent (Denville Scientific, #E2400).

### *In vitro* reconstitution of Pol IV-mediated TLS

Rolling circle templates were constructed as described previously (1, 2). Pol IV-mediated TLS was reconstituted within the *E. coli* replisome as described previously (1). Briefly, a lesion-containing rolling circle template (1 nM) was mixed with DnaB_6_ (50 nM), τ_3_δδ’χψ (20 nM), β_2_ clamp (20 nM), Pol III core (60 nM), ATPγS (50 μM), dCTP/dGTP (each 60 μM) in 50 mM Hepes-KOH (pH8) on ice and incubated at 37 °C for 6 min. Then, replication was initiated by the addition of a pre-warmed 10 X mixture of SSB_4_ (500 nM), DnaG (100 nM), ATP (1 mM), CTP/GTP/UTP (each 250 μM), [α-^32^P]-dATP. 10 seconds after the addition of the initiation mixture, a mock or Pol IV was added to the reactions and the reactions continued at 37 °C for 12 min. Reactions were quenched by the addition of EDTA (24 mM). Concentrations are all final in the replication reactions. Replication products were resolved in a 0.6% denaturing alkaline agarose gel and visualized by autoradiography. From the autoradiograph, relative abundance of the replication products at the resolution limit (RL) was quantified using ImageJ.

### Measuring the interaction between Pol IV and β_2_ clamp by biolayer interferometry

Purified PK-His6-Pol IV^LF^ and its variants were immobilized onto Octet NTA biosensors (Sartorius, 18-5103) and incubated with varying concentrations of β_2_ clamp (31.3 to 8,000 nM as a dimer). Immobilization and binding reactions took place in binding buffer (50 mM Hepes-NaOH, pH7.5/100 mM NaCl/5 mM β-mercaptoethanol/0.05% Tween20). Shifts in the interference pattern upon interaction between Pol IV^LF^ and β_2_ clamp were measured in real-time using Octet RED384 (Sartorius). To determine equilibrium dissociation constants, a one site total binding model was fitted to the background (responses to β_2_ clamp in the absence of immobilized Pol IV^LF^)-subtracted steady-state responses at varying concentrations of β_2_ clamp. One site total binding model: R = Rmax × A / (K_D_ + A) + NS × A + Background. R, response; Rmax, maximum response; A, concentration of analyte; NS, slope of nonspecific binding; Background, response in the absence of analytes.

### Measuring the damage-induced SOS responses

LB medium was inoculated with SOS reporter strains and incubated at 37 °C overnight with aeration. The overnight cultures were diluted in fresh LB to OD_600_ 0.1 and incubated at 37 °C with aeration until OD_600_ reached between 0.2 and 0.3. These cultures were treated with NFZ (60 μM, dissolved in dimethyl formamide) or MMS (7.5 mM) and further incubated at 37 °C with aeration for 1 hour. Then, cells were collected by centrifugation and washed with ice-cold phosphate buffered saline (PBS) twice. Washed cells were fixed with PBS containing 4% formaldehyde. Fixed cells were washed with ice-cold PBS twice and resuspended in ice-cold PBS. Fluorescence signals from individual cells were measured using a flow cytometer (BD Accuri C6). Means and standard deviations of fluorescence signals from about one hundred thousand cells were determined.

### Measuring pathway choice in cells

Construction of control and lesion-containing plasmids: Plasmids carrying a single lesion for genomic integration were constructed as described previously (8). Briefly, 3meA-containing and its lesion-free control plasmids were constructed by inserting an oligomer (5′-GCAAGTT<u>A</u>ACACG-3′, <u>A</u>, 3-deaza-methyl dA (lesion) or dA (control)) into a gapped-duplex pGP1/2 (amp^R^), creating an in frame *lacZ* gene. N^2^-furfuryl dG-containing and its lesion-free control plasmids were similarly created with lesion-containing and control oligomers (5’-CTACCT<u>G</u>TGGACGGCTGCGA-3’, <u>G</u>, N^2^-furfuryl dG (lesion) or dG (control)) as described previously (2).

In cell pathway choice assays: Assays were performed as described previously (8). Briefly, a control or lesion-containing plasmid was introduced into the recipient strains together with a transformation control plasmid (pVP146, tet^R^) by electroporation using GenePulser Xcell (BioRad). These electroporated cells were first resuspended in 1 ml of super optimal broth with catabolic repressor (SOC), then 500 μl of resuspended cells was transferred into 2 ml of LB containing 0.2 mM IPTG to induce the expression of integrase and excisionase. These cells were further incubated at 37 °C for 45 minutes. A portion of these cells were plated on LB containing 10 μg/ml tetracycline to measure the transformation efficiency, and the rest of the cells were plated on LB containing ampicillin (50 μg/ml) and X-gal (80 μg/ml) to select for integrants (amp^R^) and monitor TLS among the integrants (*lacZ^+^* phenotype, formation of sectored blue/white colonies in the presence of X-gal). After 24-hour incubation at 37 °C, colonies were counted, and choices of DA (white colony counts/total colony counts) and TLS (sectored blue-white colony counts/total colony counts) were determined.

### Imaging

Samples were prepared for microscopy and imaged exactly as described previously (4). In brief, imaging cultures were grown in supplemented M9 medium with 0.4% glucose as the carbon source. IPTG was added to a final concentration of 0.5 mM to induce expression of the SSB-mYPet fusion. When cultures reached OD_600_ ≈ 0.15, an aliquot was harvested by centrifugation and deposited on an agarose pad containing 100 mM MMS, which was sandwiched between cleaned coverslips. The sample was incubated on the pad for 20 min before imaging.

Imaging was performed in a custom-built fluorescence microscope described previously (4). A Nikon TE2000 microscope was equipped with a Nikon CFI Apo ×100/1.49 NA TIRF objective, dichroic mirrors and bandpass filters (Chroma), and a Hamamatsu ImageEM C9100-13 EMCCD camera. Excitation was provided by 514 nm and 561 nm lasers (Coherent Sapphire) and PAmCherry photoactivation was achieved using a 405 nm laser (Coherent OBIS). Lasers were focused to the objective back focal plane to achieve highly inclined thin illumination (16). Computer-controlled shutters were used to automate excitation sequences and the 405 nm photoactivation power was adjusted to control the density of photoactivated Pol IV-PAmCherry molecules. For all experiments, at least three different biological replicates (imaging cultures) were imaged on at least two different days.

Analysis of microscopy data was performed exactly as described previously (4). In brief, brightfield images of cells were segmented using the MicrobeTracker package (17) and detection and tracking of Pol IV-PAmCherry molecules was performed using the u-track package (18, 19). Pol IV-PAmCherry molecules were fit to 2D Gaussian approximations of the point spread function (PSF) and static molecules were identified based on their PSF width. For analysis of SSB-mYPet foci, the first five frames of 514 nm excitation were averaged and then spot detection was performed using u-track. To remove a small number of poorly fit weak fluorescent spots, foci with background values less than the camera offset level (1,500 counts) were removed. To avoid possible crosstalk from the SSB-mYPet foci, the small fraction of cells containing a Pol IV-PAmCherry localization in the first frame of PALM excitation were excluded from the dataset.

Colocalization between Pol IV-PAmCherry molecules and SSB-mYPet foci was analyzed using radial distribution function analysis exactly as described previously (4). The distance between each Pol IV-PAmCherry trajectory and the nearest SSB-mYPet focus in the cell was calculated and aggregated across all cells, giving an experimental distance distribution. Subsequently, Pol IV-PAmCherry localizations were randomly simulated in the cells and a random distance distribution was generated. The experimental distance distribution was normalized by the simulated one to give the radial distribution function *g*(*r*) for all distance values *r*. This procedure was repeated for 100 simulated distance distributions and the resulting *g*(*r*) curves were averaged to give the final average *g*(*r*) curve. To validate the analysis, a separate random distance distribution was generated, and the same procedure was followed to generate an average random *g*(*r*) curve; this random *g*(*r*) curve should be approximately equal to one at all distances.

